# Deep learning neural network prediction method improves proteome profiling of vascular sap of grapevines during Pierce’s disease development

**DOI:** 10.1101/2020.07.18.210153

**Authors:** Cíntia H. D. Sagawa, Paulo A. Zaini, Renata de A. B. Assis, Houston Saxe, Michelle Salemi, Aaron Jacobson, Brett S. Phinney, Abhaya M. Dandekar

## Abstract

Plant secretome studies have shown the importance of plant defense proteins in the vascular system against pathogens. Studies on Pierce’s disease of grapevines caused by the xylem-limited bacteria *Xylella fastidiosa* (*Xf*) have detected proteins and pathways associated to its pathobiology. Despite the biological importance of the secreted proteins in the extracellular space to plant survival and development, proteome studies are scarce due to technical and technological challenges. Deep learning neural network prediction methods can provide powerful tools for improving proteome profiling by data-independent acquisition (DIA). We aimed to explore the potential of this strategy by combining it with *in silico* spectral library prediction tool, Prosit, to analyze the proteome of vascular leaf sap of grapevines with Pierce’s disease. The results demonstrate that the combination of DIA and Prosit increased the total number of identified proteins from 145 to 360 for grapevines and 18 to 90 for *Xf*. The new proteins increased the range of molecular weight, assisted on the identification of more exclusive peptides per protein, and increased the identification of low abundance proteins. These increases allowed the identification of new functional pathways associated with cellular responses to oxidative stress to be further investigated.

## 1. Introduction

The vascular system is essential for the exchange of information and resource allocation throughout the plant, from roots to aerial tissues. It is composed of two types of vascular tissues: phloem and xylem. The phloem sap contains photoassimilates and other macromolecules that move throughout the plant from areas of synthesis or excess (source) to areas of use (sink) and storage [1]. The xylem sap transports water and nutrients from roots to aerial tissues, driven by a difference in water potential due to transpiration (Tanner and Beevers, 2001). Recent studies have shown that the xylem can also contain a wide range of proteins involved in various biological processes involved in growth regulation, protection against environmental stress, homeostasis, gas exchanges, cell to cell adhesions, and plant defense against pathogens [3]. These processes are dependent on vesicular trafficking of proteins to the extracellular space, which can either follow conventional or unconventional secretion routes in plant cells. The conventional secretion in plants requires signal peptides in the N-terminus or proper recognition signals to direct them to the endomembrane system pathway, while proteins that follow the unconventional secretion route lack these signals [4]. Plant secretome studies have shown that proteins that follow unconventional secretion can allow plants to respond to a wider range of extracellular stresses and stimuli, facilitating defense responses under stress [4], [5]. Despite the biological importance of the secreted proteins in the extracellular space to plant survival and development, proteome studies are scarce due to technical and technological challenges.

Studies on the role of vascular sap have helped to better understand plant responses to vascular plant diseases (Yadeta and Thomma, 2013). The Gram-negative gammaproteobacteria *Xylella fastidiosa* (*Xf*) is a xylem-limited pathogen that colonizes several economically important crops worldwide causing deadly diseases such as Pierce’s disease in grapevines (PD) (Davis et al., 1978), Citrus Variegated Chlorosis (CVC) [8] and most recently Olive Quick Decline Syndrome (OQDS) in Europe (Martelli, 2016). Due to the significant economic impact on the production of citrus in Brazil, *X. fastidiosa* was the first plant pathogen to have its genome sequence determined [10]. The genomic landscape provided an initial description of potential virulence factors and revealed the absence of a type III secretion system commonly employed by plant pathogens to deliver virulence effectors inside plant cells. Molecular and cellular studies followed proposing that the mechanism of disease symptoms would be associated with biofilm formation and xylem blockage triggering the observed disease symptoms [11]–[15]. Additionally, genomics and proteomics have shown the importance of virulence factors secreted by the type II secretion system and outer membrane vesicles for symptom development (Nascimento et al. 2016; Gouran et al. 2016; Santiago et al. 2016; Cianciotto and White 2017; Feitosa-Junior et al. 2019). These studies highlighted the molecular complexity of the plant-pathogen interaction that takes place in the vascular system.

The first study on xylem sap proteomics in grapevines was performed in sap bleedings from the cultivar Chardonnay (Agüero et al. 2008), which revealed only ten proteins from two-dimensional (2D) gel electrophoresis analysis. As new technologies and proteomic approaches became more sensitive, more proteins were found in the vascular sap of grapevines, increasing the number of identified proteins to 200 varying from 20 to 75 kDa showing differences among resistant and susceptible cultivars to PD (Delaunois et al. 2013). The importance of proteins in the plant response to *X. fastidiosa* was initially shown by Yang et al. (2011) in a proteomic study of stems from infected grapevines. This study revealed thaumatin-like, pathogenesis-related protein 10 and three heat shock proteins were significantly overexpressed in PD-resistant varieties of grapes (Yang et al. 2011). Another study also conducted on the stem of infected grapevines of PD-tolerant and susceptible cultivars identified more than 200 proteins associated with disease resistance, energy metabolism, protein processing and degradation, biosynthesis, stress-related functions, cell wall biogenesis, signal transduction, and ROS detoxification among others [23]. The most recent published study conducted on sap bleeding of infected grapevines highlighted 91 proteins. The novelty of this study was the incorporation of structural data into the proteomic data analysis to enhance the identification of functionally relevant protein candidates that would not be detected from simple amino acid sequence alignments. This study highlighted pathogenesis-related proteins, chitinases, and β-1, 3-glucanases as crucial players in the defense against *X. fastidiosa* [25]. These studies greatly enhanced our understanding of xylem sap physiology; however, they were restricted to more abundant proteins which we have learned to be only a small fraction of xylem sap complexity.

The standard approach in proteomic studies was 2D gel electrophoresis for many years due to its robustness and compatibility with bottom-up (shotgun) proteomics in which the crude protein extract is digested directly for analysis. However, the limitations regarding reproducibility and narrow dynamic range of high abundance proteins masked low abundant counterpart, limiting those analyses [26]. Electrophoresis gels can now be replaced by liquid chromatography coupled with tandem mass spectrometry (LC-MS/MS), which has become the most used method to measure the different states and abundance of proteins, lipids and other metabolites [27].

One of the acquisition schemes of tandem mass spectrometry is called data-independent acquisition (DIA) which is based on the acquisition of fragment-ion information for all precursor ions until the desired mass range has been covered, as demonstrated by the sequential window acquisition of all the theoretical mass spectra (SWATH) approach [28]. DIA has been used to identify and quantify thousands of proteins without performing fractionation, increasing reproducibility, and requiring a small amount of protein [27], [29], [30]. Although it improves protein detection with higher reproducibility, the lack of accurate predictive models for fragment ion intensities has impaired its full potential. DIA analysis often uses peptide physiochemical properties stored in spectral libraries or chromatogram libraries. These properties can include information on peptide retention time, product ion m/z, product ion intensity and ion mobility among others [31], [32]. Using this information can ensure confident peptide identification and quantification. Two methods exist to obtain this information, one is experimental and the other is predictive. An example of a predictive method is the deep learning architecture termed Prosit which was created to take advantage of a large number of synthetic peptides and tandem mass spectra generated within the ProteomeTools project to predict with high quality both chromatographic retention time and fragment ion intensity of any peptide [33]. Here we demonstrate the improved performance of integrating Prosit into the DIA pipeline. By reanalyzing our DIA data of the vascular leaf sap of grapevines infected by *X. fastidiosa* compared with healthy plants, we increased the number of identified proteins depicting a deeper description of this plant pathogen interface and generated spectral libraries for DIA analysis of *Vitis vinifera* and *Xylella fastidiosa* that can be incorporated in future proteome studies.

## 2. Material and methods

### 2.1. Plant material and *X. fastidiosa* inoculation

Clonal grapevine plants (*Vitis vinifera* L. cv. ’Thompson Seedless’) were generated from cuttings using green canes from the current season’s growth. Each cutting was approximately 6 inches long and contained two nodes, with a petiole originating from the top node that supported approximately one square inch of leaf area to maintain minimal photosynthesis during rooting. These prepared cuttings were placed into an EZ-Clone aeroponic cloning system that circulates water purified by reverse osmosis. Roots begin to self-generate after two weeks, and the rooted cuttings were potted after three-weeks and grown in a greenhouse. New plant growths was trained to a single cane by removing any lateral shoots that emerged. The single cane plants were topped at the height of 1 meter, and additional lateral shoots were removed as they emerged during the experiment. After ten-weeks, the grapevines were infected at 8–12 cm above soil level by punching with a needle gauge to inoculate 20 μL of cultured cells of *Xylella fastidiosa* Temecula1 (*Xf*; ATCC 700964) into the stem as described by Nascimento et al. (2016). The bacterial culture was grown on PD3 medium at 2×10^8^ cells/mL incubated with aeration (120 rpm) at 28°C. After inoculation, plants were placed in the greenhouse in a randomized block design and monitored for 12 weeks post inoculation until leaf symptoms developed.

### 2.3. Vascular sap extraction and *X. fastidiosa* quantification

Vascular leaf sap was collected from ten leaves above the inoculation point using a pressure chamber (Soil Moisture Equipment Corp., Santa Barbra, CA, USA). Pressure was applied to each leaf blade and the sap collected from the end of the petiole. The leaf blade was placed inside the pressurized chamber leaving only the cut surface of the petiole exposed to release the vascular content, which was collected using a micropipette and stored in a tube on ice during harvest. Pools of about ten leaves above the inoculation point from one plant made one sample (500 uL - 1000 uL). Before processing with the sample preparation for proteomics analysis, an aliquot of 25 uL was reserved from each sample for extraction of DNA with MasterPure™ kit (Epicentre) and bacterial cell count was measured using qPCR (TaqMan™). The primers used were HL5 and HL6 described by Francis et al. (2006). A standard curve was used based on a known serial dilution of *Xf* cells measured by OD_600_.

### 2.4. Protein digestion of vascular leaf sap

Up to one milliliter of vascular leaf sap was collected from each plant (pooled from 10 leaves) and a total of three plants per group (Healthy and Diseased) were used. Samples were centrifuged at 5,000 rcf for 5 min at 4°C. The supernatant containing the vascular leaf sap was transferred to a new tube. Total protein content was quantified by Qubit™ Protein Assay Kit (Thermo Fisher Scientific). Sap containing 100 ug of protein was freeze-dried and resuspended in 5% SDS and 50mM triethylammonium bicarbonate (TEAB) at pH 7.55 to a concentration of 0.5 ug/uL. Digestions with trypsin followed the S-Trap™ Micro Spin Column Digestion Protocol with few modifications. Initially, 10 mM dithiothreitol (DTT) was added and incubated at 50°C for 10 min and rested at room temperature for 10 min. Next, 5 mM iodoacetamide (IAA) was added and incubated at room temperature for 30 min in the dark. The samples were acidified with 12% phosphoric acid followed by the addition of 2.348 mL of freshly made S-trap buffer (90% methanol, 100 mM TEAB, pH 7.1) and mixed immediately by inversion. The entire acidified lysate/St-buffer mix was transferred to the S-trap spin column (650 uL at a time) and centrifuged at 3,000 rcf for 1 min or until all the solution passed through the column. Columns were washed with 400 uL of S-trap buffer and centrifuged at 4,000 rcf until dry. Columns were transferred to a clean elution tube. Trypsin enzyme digest buffer was carefully added (1:25 enzyme: total protein in 121 uL 50mM TEAB, pH 8.0) to the column and followed by incubation at 37°C overnight. After the first hour, the trypsin digestion step was repeated. Peptide elution steps included 80 uL of 50 mM TEAB (pH 8.0) followed by centrifugation at 1,000 rcf for 1 min, 80 uL of 0.5% formic acid followed by centrifugation at 1,000 rcf for 1 min, 80 uL of the solution containing 50% acetonitrile and 0.5% formic acid followed by centrifugation at 4,000 rcf for 1 min. The final pooled elution was dried down in a speed-vacuum. Peptides were resuspended in 0.1% TFA 2% ACN and quantified using Pierce™ Quantitative Fluorometric Peptide Assay (Thermo Fisher Scientific). Equal portions of all samples were mixed together to make a reference sample to be run multiple times for chromatogram library runs.

### 2.5. Liquid chromatography tandem mass spectrometry

The next steps were processed at the UC Davis Proteomics Core Facility. Peptides were trapped on a Thermo PepMap trap and separated on an Easy-spray 100 um × 25 cm C18 column using a Dionex Ultimate 3000 nUPLC at 200 nl/min. Solvent A= 0.1% formic acid, Solvent B = 100% Acetonitrile 0.1% formic acid. Gradient conditions = 2%B to 50%B over 60 minutes, followed by a 50%-99% B in 6 minutes and then held for 3 minutes than 99%B to 2%B in 2 minutes and total run time of 90 minutes using Thermo Scientific Fusion Lumos mass spectrometer running in Data Independent Acquisition (DIA) mode.

### 2.6. Chromatogram library creation

Six-gas phase fractionated (GFP) chromatogram library injections were made using staggered 4 Da isolation widows. GFP1 = 400-500 m/z, GFP2 = 500-600 m/z, GFP3 = 600-700 m/z, GFP4 = 700-800 m/z, GFP5 = 800-900 m/z, GFP6 = 900-1000 m/z, mass spectra were acquired using a collision energy of 35, resolution of 30 K, maximum inject time of 54 ms and a AGC target of 50K. Each individual sample was run in DIA mode with staggered isolation windows of 12 Da in the range 400-1000 m/z.

### 2.7. Analytic samples, data analysis and raw data processing

Each individual sample was run in DIA mode using the same settings as the chromatogram library runs except using staggered isolation windows of 12 Da in the m/z range 400-1000 m/z. DIA data was analyzed using Scaffold DIA v.2.0.0 (Proteome Software, Portland, OR, USA). Raw data files were converted to mzML format using ProteoWizard v.3.0.11748 [35].

### 2.8. Spectral library search

The Reference Spectral Library was created by EncyclopeDIA v.0.9.2. Chromatogram library samples were individually searched against Prosit predicted databases created using Prosit online server (https://www.proteomicsdb.org/prosit/) and converted for ScaffoldDIA using the Encyclopedia tools [32]. The input for the Prosit prediction consisted of UniProt proteome UP000009183 (*Vitis vinifera*, Grape), UniProt proteome UP000000812 (*Xylella fastidiosa*) and 114 common laboratory contaminants (https://www.thegpm.org/crap/) with a peptide mass tolerance of 10.0 ppm and a fragment mass tolerance of 10.0 ppm. Variable modifications considered were oxidation of methionine and static modifications were carbamidomethyl of cysteine. The digestion enzyme was assumed to be Trypsin with a maximum of 1 missed cleavage site(s) allowed. Only peptides with charges in the range [2–3] and length in the range [6–30] were considered. Peptides identified in each search were filtered by Percolator (3.01.nightly-13-655e4c7-dirty) [36]–[38] to achieve a maximum FDR of 0.01. Individual search results were combined, and peptides were again filtered to an FDR threshold of 0.01 for inclusion in the reference library. A summary of the workflow is presented in Figure 1.

**Figure 1.**
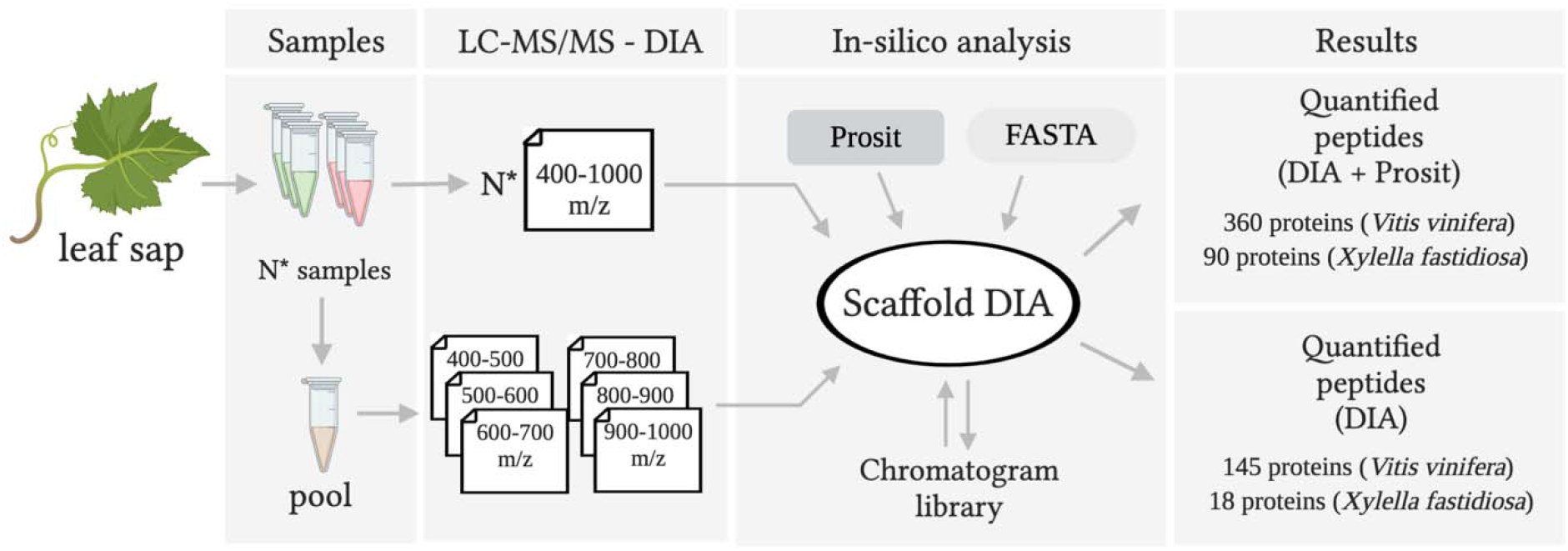
Quantification of peptides with chromatogram libraries workflow. The chromatogram library generation was based on Searle et al. (2018). In summary, each quantitative replicate (analytic samples) for each group was measured by wide-window DIA experiment (400-1000 m/z) besides the collection of several staggered narrow-window DIA experiment from the pooled sample of all samples. Afterwards, these narrow-window experiments have 2 m/z precursor isolation targeting every peptide between 400 and 1000 m/z. The peptides anchors were detected using ScaffoldDIA. Chromatographic data about each peptide was stored in a chromatogram library with retention times, peak shape, fragment ion intensities, and known interferences tuned specifically for the LC-MS/MS setting. ScaffoldDIA uses these precise coordinates for m/z, time, and intensity to detect peptides in the quantitative samples generating the DIA results box. Alternatively, in addition to the chromatogram library generated, a predicted library created using Prosit and FASTA information was added to determine quantified peptides and generated the DIA + Prosit results box. Created with BioRender.com.

### 2.9. Quantification and criteria for protein identification

Peptide quantification was performed by EncyclopeDIA v. 0.9.2. For each peptide, the five highest quality fragment ions were selected for quantitation. Proteins that contained similar peptides and could not be differentiated based on MS/MS analysis were grouped to satisfy the principles of parsimony. Proteins with a minimum of 2 identified peptides were thresholder to achieve a protein FDR threshold of 1.0%.

### 2.10. Functional enrichment analysis

The functional analysis of proteomics of vascular leaf sap of grapevines was performed by the online software Metascape [39] using the express analysis settings. The up and downregulated *Vitis vinifera* protein IDs of diseased samples were converted into the corresponding *Arabidopsis* homolog protein IDs and analyzed independently. The *Arabidopsis* homologs were identified in TAIR using Protein Basic Local Alignment Search Tool (BLASTP). Metascape identified pathways and process enrichment analysis defined by the Kyoto Encyclopedia of Genes and Genomes (KEGG). P-value was adjusted by the method of Benjamin-Hochberg to control the false discovery rate (FDR).

## 3. Results

### 3.1. Creating a DIA library and improving the datamining of xylem proteome data

In this study, we compared the proteome of vascular leaf sap from healthy grapevines to those developing PD symptoms due to *X. fastidiosa* (*Xf*) infection. Infection was confirmed by qPCR that quantified a high number of bacterial cells 1.5×10^9^ cells/mL present in the diseased samples (Table S1). The vascular system is particularly crucial for this pathosystem as *Xf* cells are restricted to this microenvironment within plants. Thus, much of its interaction with the host occurs on the surface of xylem cells. As proteomic methods and equipment are rapidly evolving, we investigated the effect of a new deep neural proteome prediction method, Prosit, to identify proteins from mass spec data applied on Data Independent Acquisition (DIA) currently in use.

Figure 2a shows the proteomics results from vascular leaf sap of grapevines *Vitis vinifera* (VIT) at 12 weeks post-inoculation with *Xf*. DIA analysis identified 145 and 18 proteins for VIT (Table S2) and *Xf* (Table S3), respectively. After integrating Prosit into database search pipelines, the number of proteins increased by more than 148% for VIT and 400% for *Xf*, to a final total of 360 and 90 proteins (Tables S4 and S5). Only six VIT proteins were identified exclusively without Prosit and 221 only by integrating Prosit (DIA+Prosit), with 139 detected in either approach for VIT (Fig.2b, Table S6). Among the six VIT proteins identified by DIA only, four are peroxidases (VIT_01s0010g01950, VIT_01s0010g01960, VIT_01s0010g02000, VIT_01s0010g02010), an uncharacterized protein with serine-type endopeptidase activity (VIT_16s0098g01160), and a Glyco_hydro_18 domain-containing protein (VIT_16s0050g02220). Nevertheless, the proteins detected exclusively by Prosit were associated with many more molecular functions, including cell adhesion molecules, scaffold/adaptors proteins, chaperones, translational proteins, transporters, and nucleic acid-binding proteins. Regarding the *Xf* bacterial proteins, 18 proteins were identified by both methods; however, DIA+Prosit allowed the detection of an additional 72 proteins that were not present in the DIA data (Table S7).

**Figure 2.**
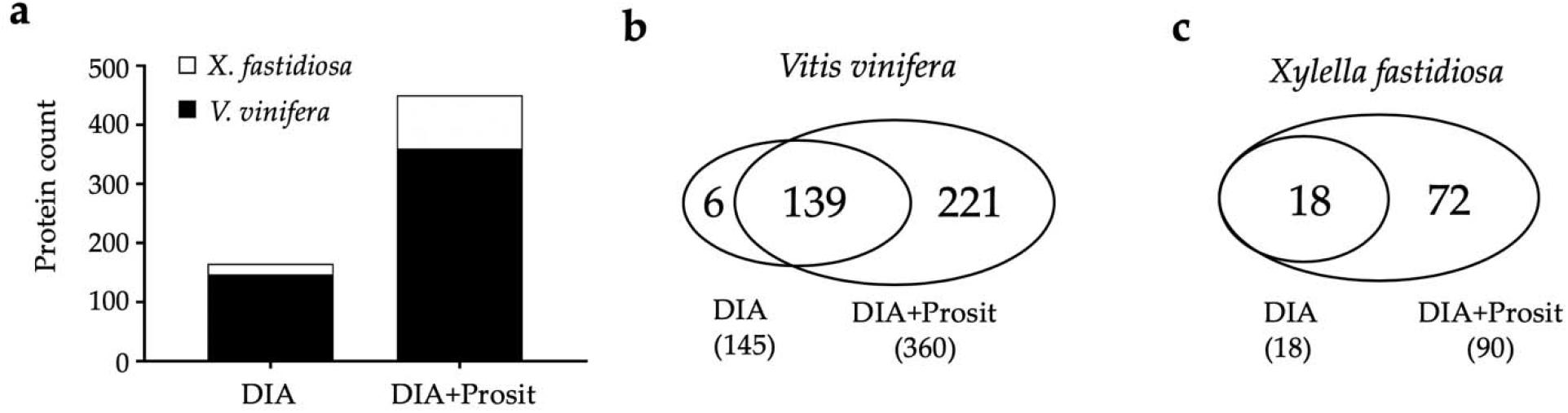
Proteomic analysis of *Vitis vinifera* and *Xylella fastidiosa*: a) total proteins identified by data-independent acquisition (DIA) and DIA+Prosit; b) Venn diagram of the number of proteins identified by each method for *V. vinifera*; and c) for *X. fastidiosa*.

The application of Prosit to our data substantially increased the number of proteins with a molecular weight below 100 kDa. The range of molecular weight varied from 12 kDa to 217 kDa in DIA data and 8 kDa to 217 kDa in DIA+Prosit data. A breakdown of identified proteins on both methods by molecular weight and the number of mapped peptides is shown in Figure 3a. The smallest proteins predicted by DIA are AAI domain-containing proteins (VIT_02s0236g00020 and VIT_02s0236g00030) with 12 kDa, both upregulated in diseased plants. In addition to identifying more proteins, DIA+Prosit also increased the number of peptides identified for each protein. This is a significant advancement since we set a minimum of two mapped peptides per protein for it to be considered, considerably increasing the confidence and reducing false discoveries. The maximum of peptides identified per protein for DIA was 22, and for DIA+Prosit was 31 peptides. Most of the proteins identified after Prosit integration showed 2 to 10 peptides per protein (Fig.3b). In DIA+Prosit data, 8 kDa was the smallest protein detected, identified as BBE domain-containing protein (VIT_10s0003g05430) with a signal peptide targeting mitochondria (mTP) according to TargetP (Fig.4). Both AAI domain-containing proteins detected by DIA were also present with DIA+Prosit, and a third AAI domain-containing protein (VIT_16s0013g00070) was also detected. This is yet another important improvement as protein families with multiple members represented in a dataset gain higher scores in functional analyses such as gene ontology or pathway mapping.

**Figure 3.**
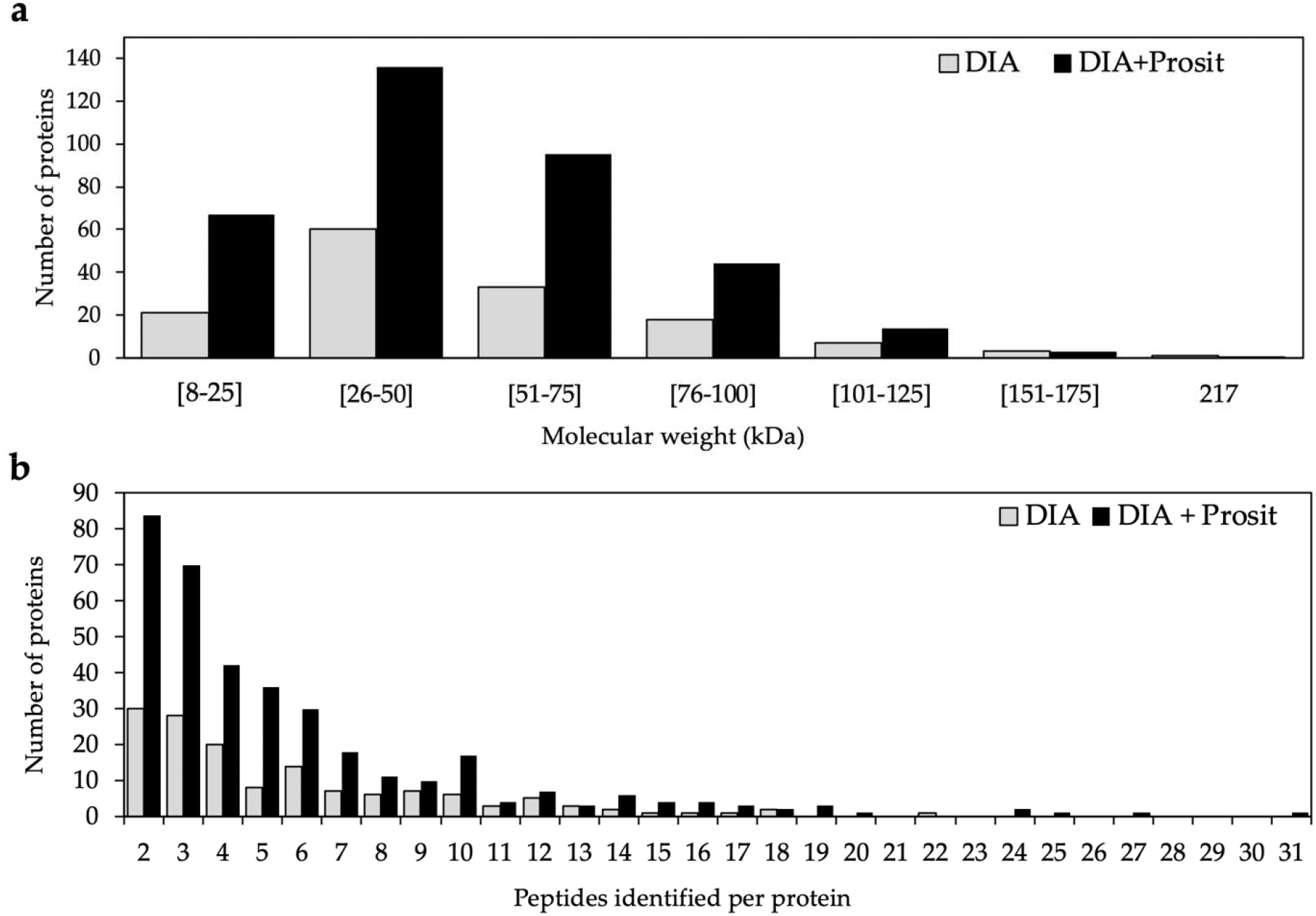
Distribution of the total number of proteins of *V. vinifera* identified by DIA and DIA+Prosit by a) molecular weight (kDa) ranging from 8 to 217 kDa. b) Identified peptide varying from 2 to 31 peptides per protein. Predicted proteins with only one peptide were discarded.

**Figure 4.**
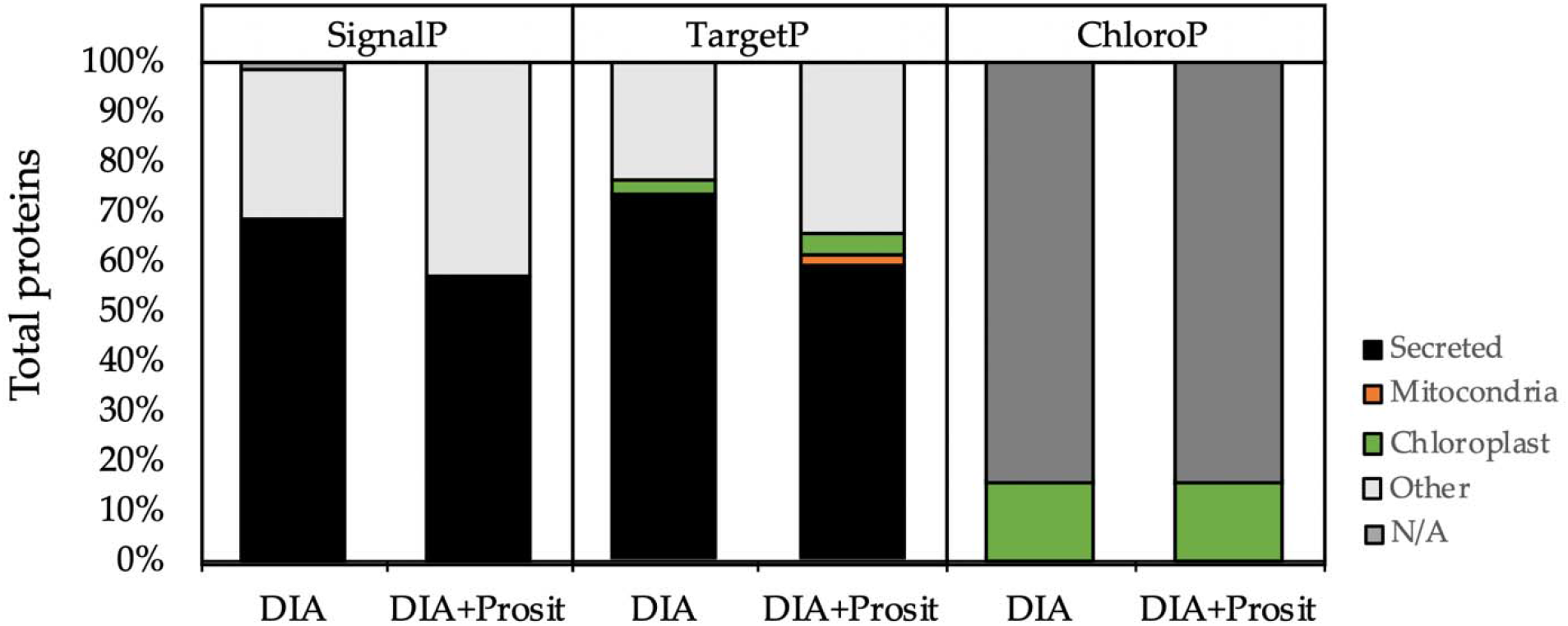
Subcellular localization prediction analysis and comparisons between DIA and DIA+Prosit data using SignalP, TargetP, and ChloroP servers. More than 50% of the total proteins identified were predicted having a signal peptide, according to SignalP and TargetP-SP. TargetP output revealed less than 3% of total proteins containing a mitochondrial targeting peptide (mTP) and less than 5% of proteins containing a chloroplast transit peptide (cTP). ChloroP predicted 16% of the collected vascular sap targeting the chloroplast by both methods. DIA considered a total of 145 proteins and DIA+Prosit, a total of 360 proteins for *V. vinifera*.

The analyzed material is an enriched vascular leaf sap; thus, we determined the proportions of proteins predicted to be secreted (Fig.4). The percentage of secreted proteins with a predicted signal peptide within the total proteins predicted for DIA was 68% (99/145), and for DIA+Prosit was 57% (205/360), according to SignalP. By using TargetP to analyze the same data sets, we found similar results: 72% and 59% for DIA and DIA+Prosit, respectively. The remaining were classified as non-secretory targeting the mitochondria (1-2%), chloroplast (3-4%), or other (23-34%). By performing the same analysis using the prediction tool ChloroP, we showed that actually, 16% of the proteins in both data sets would target chloroplasts; therefore, their presence in the xylem sap possibly reflects some degree of cellular content contamination of the samples during vacuum-assisted sap extraction or alternatively products of natural cellular and organellar degradation.

### 3.2. Regulation of proteins secreted to the xylem during Pierce’s disease

We used the MetaboAnalyst v.4.0 (https://www.metaboanalyst.ca) to visualize both proteome data sets and examined the variation between the groups and samples [40]. The variability was examined by the unsupervised principal component analysis (PCA), which showed a distinct separation between groups in both data sets, DIA and DIA+Prosit (Fig.5). In this case, the intense response to *Xf* proliferation is so marking that Prosit was unnecessary to efficiently cluster the samples by type; however, we cannot exclude the possibility Prosit would be decisive in more attenuated differences. Healthy and Diseased groups showed 85.6% variation in PC1 for DIA (Fig.5a) and 68% variation in PC1 for DIA+Prosit (Fig.5b). These results suggest the effect of *Xf* cells in the plant stress response in the proteome of the vascular leaf sap. The variation among samples explained by PC2 was 7.8% for DIA. Prosit increased the variation among samples to 16.3%, explained by PC2. For this clustering analysis, the third sample of the Healthy grapevines was discarded due to bad MS/MS data quality; therefore, a virtual sample was created using an average of the other two samples (W5 and W6; W5_W6). The protein levels in Healthy and Diseased group samples were distinct, independent of the method (Fig.S1). To further analyze the differences between methods, we analyzed the ratio-intensity of Healthy and Diseased groups and compared them to the protein abundance in both proteome data sets. The fold change of protein detection between Diseased and Healthy plants presented similar results for DIA and DIA+Prosit data (Fig.6a and 6b). However, the implementation of Prosit increased the detection of the proteins that were in low abundance, as shown by the x-axis in Figures 6a and 6b. The correlation of results obtained by both methods was significant and had an R^2^ of 0.8795 (Fig.6c), showing that the increase of protein prediction power by Prosit correlates well with the observed data without introducing bias in differential expression.

**Figure 5.**
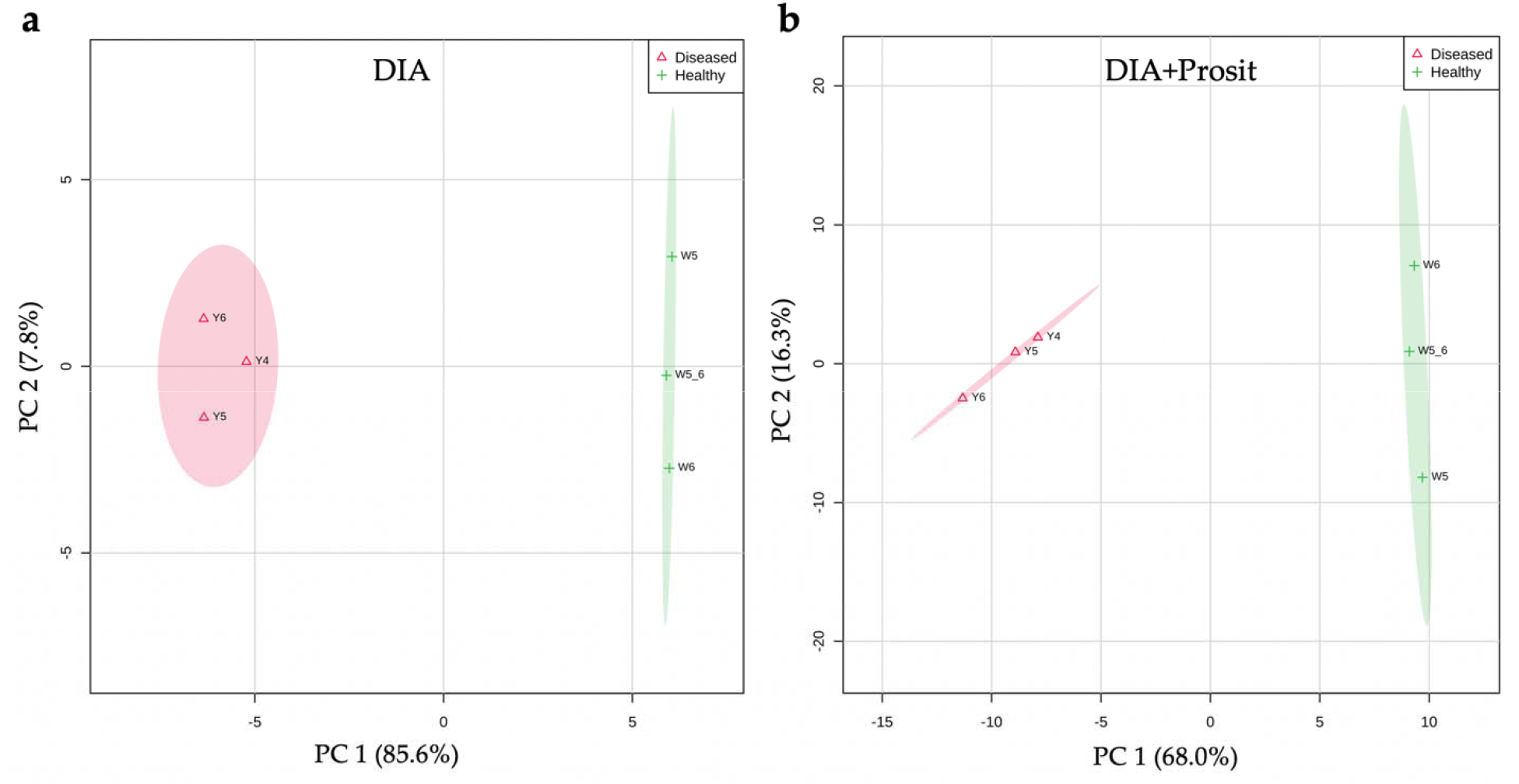
Principal Component Analysis (PCA) scores plots between PC1 and PC2 and explained variances are shown. The clear distinction between Diseased vs. Healthy proteomic data for *V. vinifera* at 12 weeks post-inoculation in both methods DIA and DIA+Prosit. W5_6 is a virtual sample made of the average of data for plants W5 and W6 (Healthy plants), and Y4, Y5, and Y5 were individual Diseased plants.

**Figure 6.**
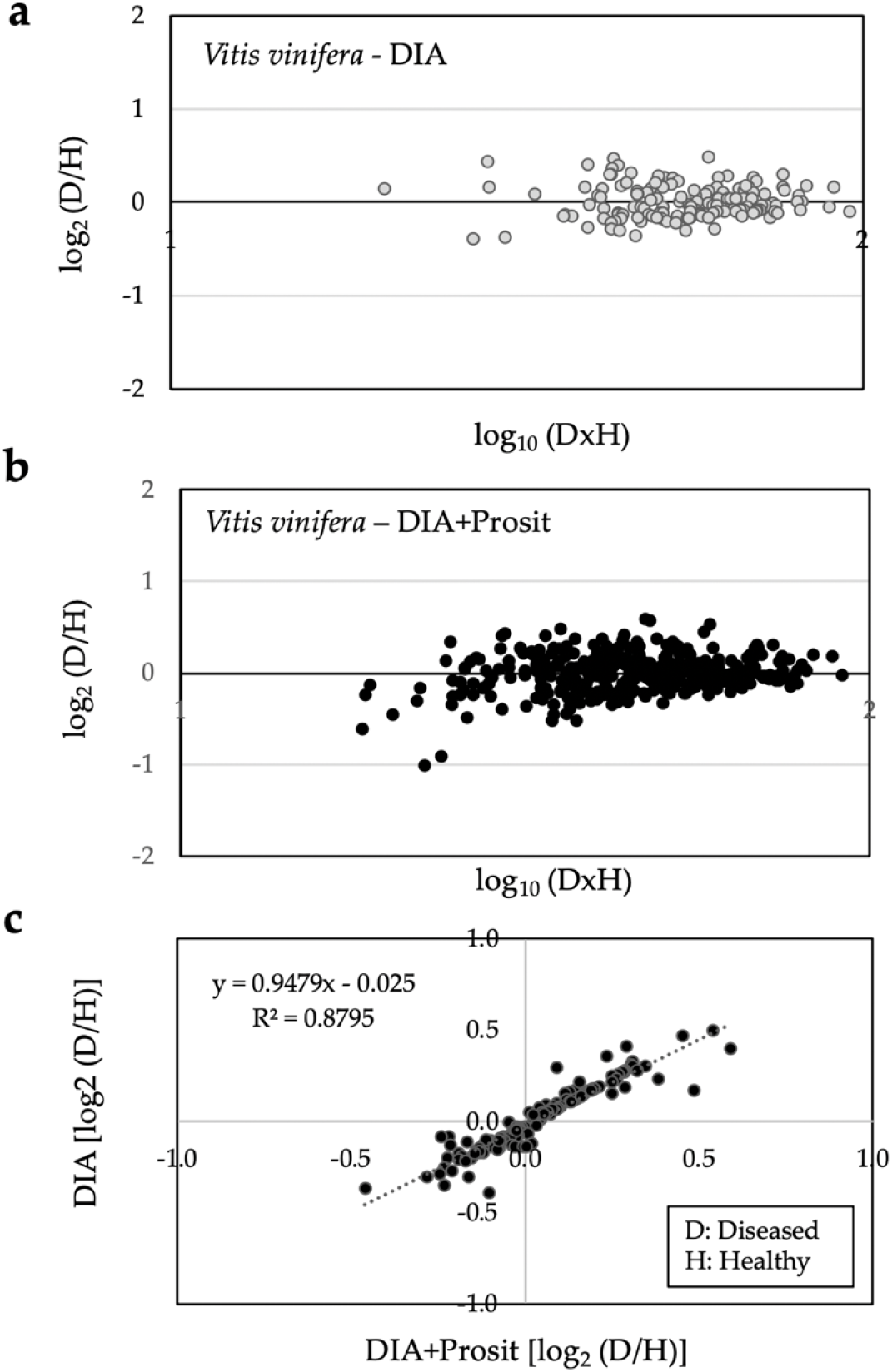
Overview of the plant response to *Xf* in Diseased samples in both data sets. Analysis of ratio-intensity plots displaying the log_2_ D/H fold-change ratio of Diseased over Healthy plants for each protein as a function of the abundance by log_10_ DxH product intensities: a) 145 proteins identified using DIA and b) 360 proteins identified by DIA+Prosit; c) Correlation between th ratios obtained from both analyses from the proteins detected in both analysis (139) with R^2^ = 0.8795 show that the incorporation of Prosit maintained provided similar results but with higher quality and expanded the detection. D: Diseased and H: Healthy plants. The log_10_ exclusive intensity data for each protein using an FDR>1% was used for both analyses.

To visualize proteins that are significantly either up or downregulated in the Diseased group, we examined volcano plots of both data sets (Fig.7). The comparison of the log_2_ fold change of the data sets and their adj. P-values by false discovery rates show a similar profile; however, the integration of Prosit allowed the identification of additional proteins that were significantly up and downregulated in the Diseased group. We observed that DIA without Prosit was more restrictive, and the maximum fold changes between Diseased and Healthy plants were not as high. The three most upregulated proteins identified by DIA+Prosit were chitinase A (VIT_16s0050g02230), Cupredoxin superfamily protein (VIT_18s0001g11180), and beta-1,3-glucanase 3 (VIT_08s0007g06060 - PR-2 family of pathogenesis-related proteins). The most downregulated proteins were Plant invertase/pectin methylesterase inhibitor superfamily (VIT_07s0005g00720), Glyco_hydro_18 domain-containing protein (VIT_06s0004g03840), and FAD-binding berberine family protein (VIT_10s0003g05470).

**Figure 7.**
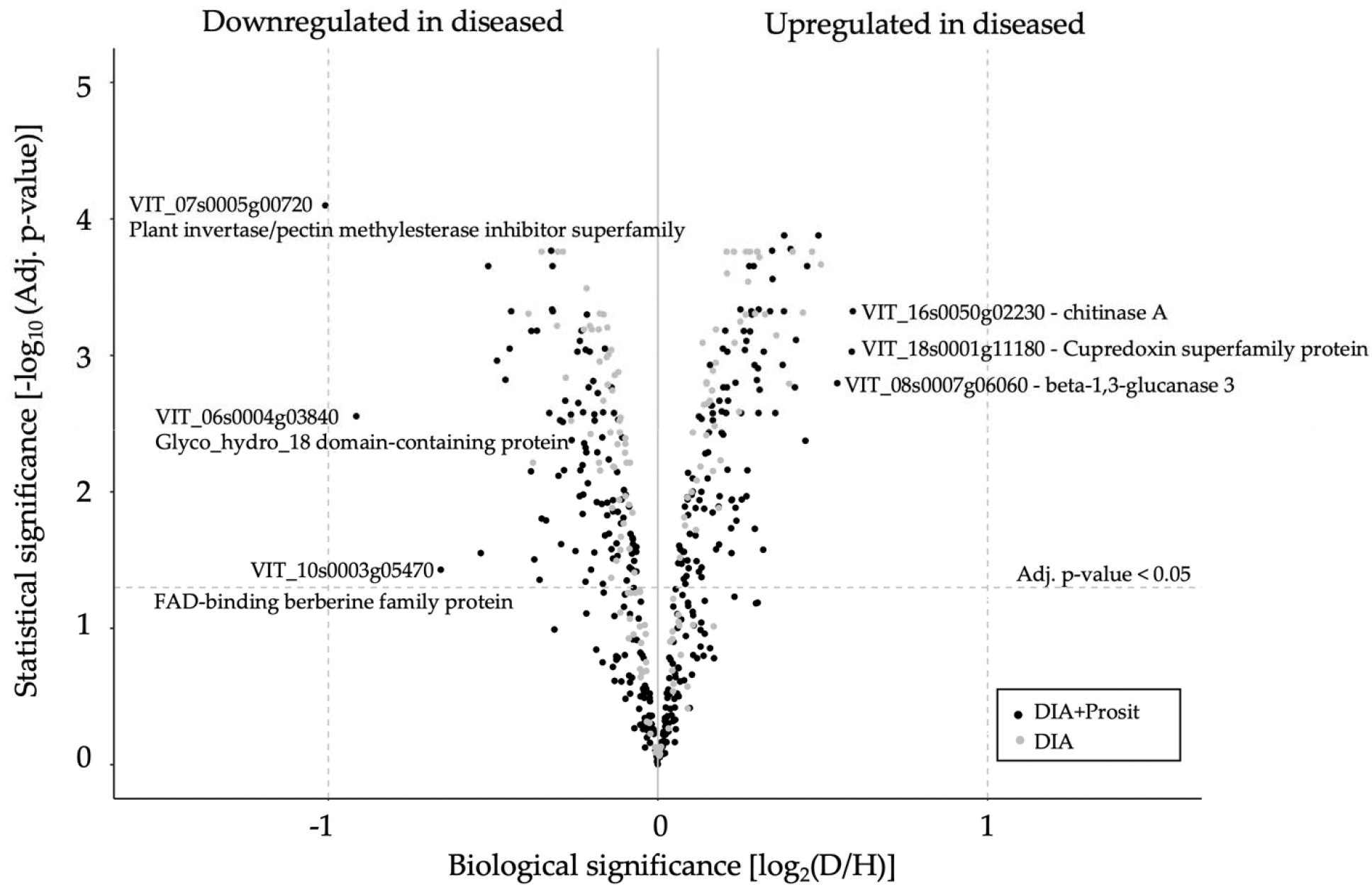
Proteome response of *V. vinifera* to *Xf* infection. Volcano plot analysis of Diseased (D) and Healthy (H) plants data identified by DIA and DIA+Prosit overlapped. Proteins identified by DIA are represented in grey dots and identified by DIA+Prosit in black dots. Adj. p-value calculated by Benjamin-Hochberg’s false discovery rate greater than or equal to 0.05 were considered significant.

For a balanced comparison between both methods, we used partial least squares - discriminant analysis (PLS-DA) of the 139 proteins that were detected by both methods. The VIP score (a metric that identifies which variables are most responsible for the differences between the classes in the analysis) was higher in DIA compared to DIA+Prosit. Among the top 25 proteins contributing to the variations among the two sample groups, we can highlight the pathogenesis-related proteins (PR1, PR2, PR3, PR4) that are upregulated in Diseased plants independent of the chosen method. Only five proteins among the top 25 in DIA were not in DIA+Prosit, and seven proteins were in DIA+Prosit and not DIA. The PLS-DA plots and the cross-validation test result are shown in Fig.S2.

### 3.3. Pathway regulation in grapevine vascular leaf sap

Representation of known enzyme pathways or protein complexes in vascular leaf sap proteome assists in the functional characterization of the plant response to infection and virulence strategies by the pathogen. The results showed that Prosit provides the identification of more pathways involved in defense during Pierce’s disease symptom development. Proteins that were up or downregulated in the Diseased group were analyzed separately to detect enriched pathways in each condition. Figures 9 and 10 show the up and downregulated proteins in both methods considering all the detected proteins for each (145 for DIA and 360 for DIA+Prosit). Most of the enriched pathways identified using DIA datasets were also present in DIA+Prosit. However, for DIA+Prosit, due to the higher number of proteins, more pathways significantly affected were revealed.

**Figure 8.**
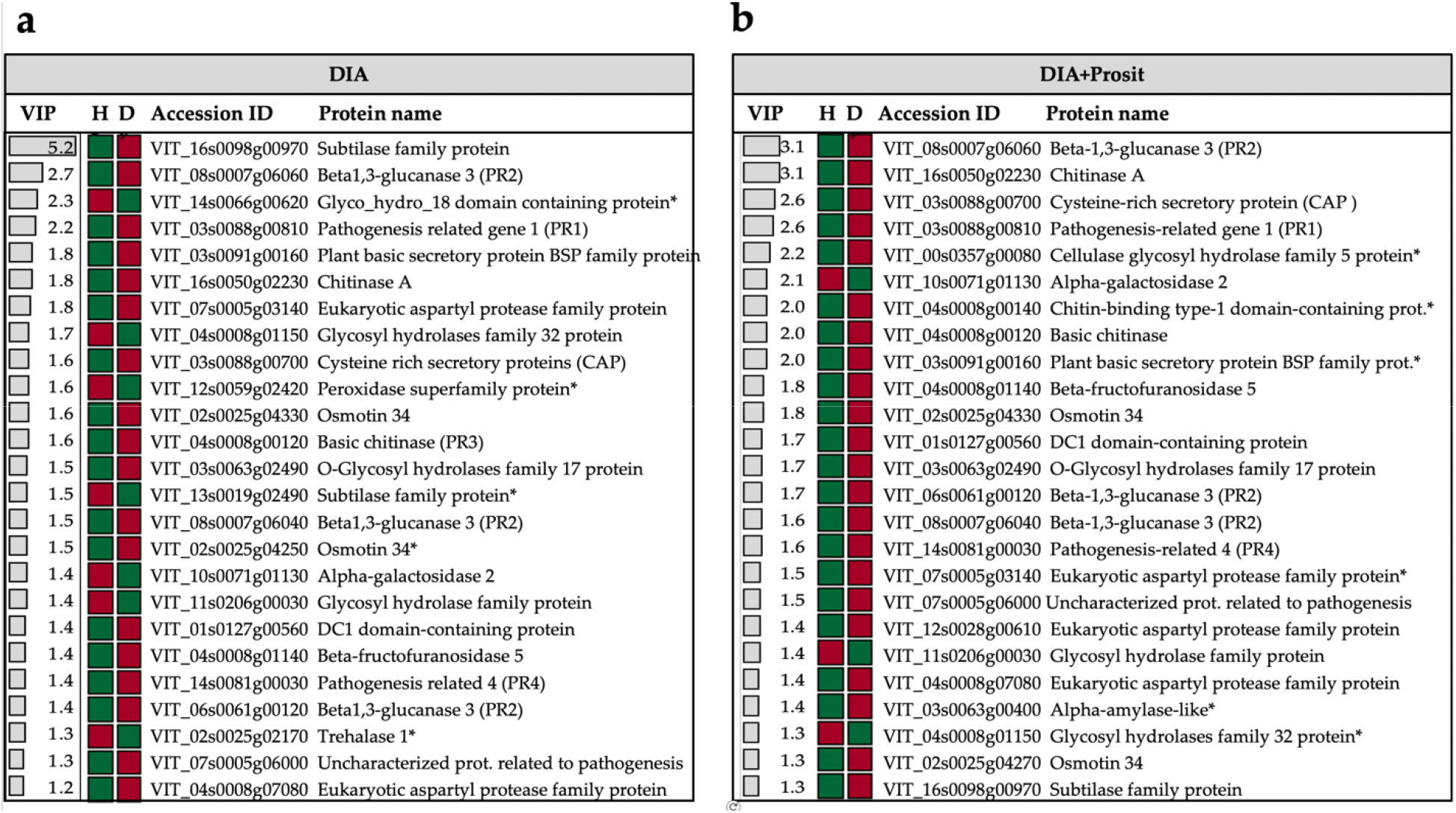
Top 25 proteins of *V. vinifera* contributing to the variance between the group observed by PLS-DA. The plot shows the variable importance in projection (VIP) scores, and the colored boxes indicate the relative intensity detected by DIA and DIA+Prosit of the corresponding protein in Diseased and Healthy plants. Red represents high and green, low exclusive intensity detected. Proteins marked with (*) are exclusive among the top 25 of the respective method.

**Figure 9.**
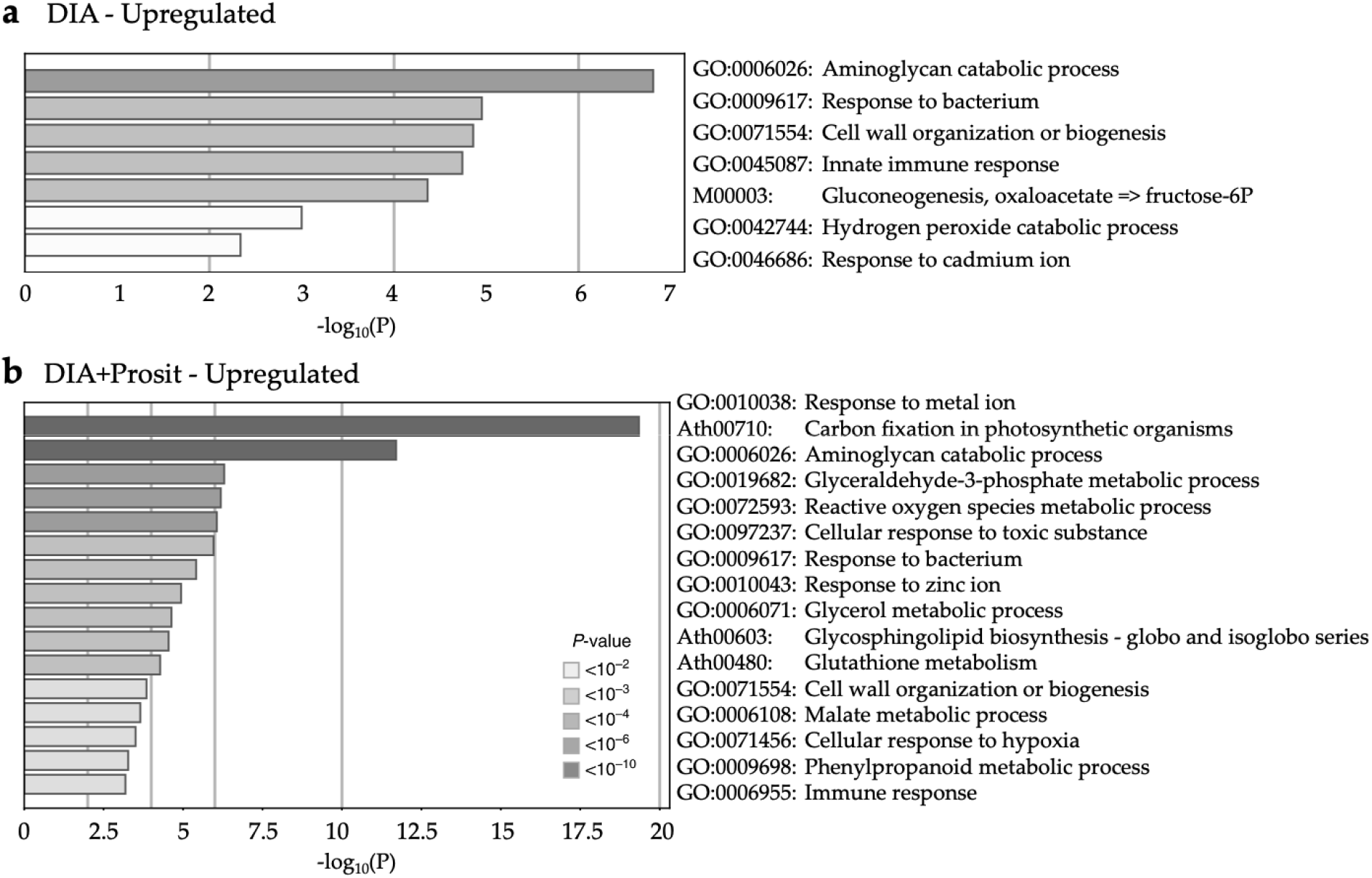
Upregulated pathways during *Xf* infection in *V. vinifera*. Non-redundant enriched ontology clusters of significantly expressed proteins upregulated during *Xf* infection (p<0.05) in a) DIA and b) DIA+Prosit data sets. DIA+Prosit allows the identification of a higher number of pathways likely involved with plant response to bacteria.

**Figure 10.**
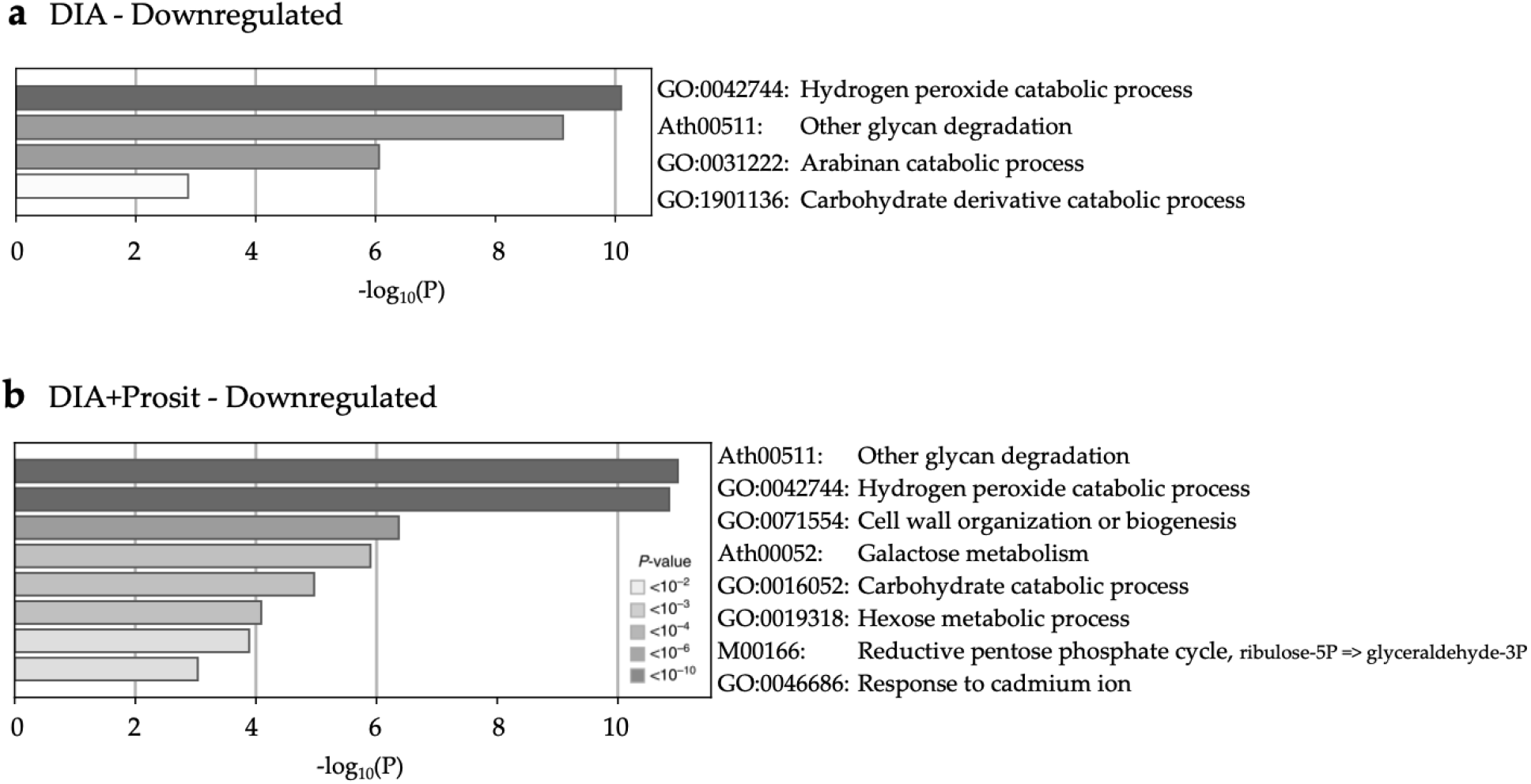
Downregulated pathways during *Xf* infection in *V. vinifera*. Non-redundant enriched ontology clusters of significantly expressed proteins downregulated during *Xf* infection (p<0.05) in a) DIA and b) DIA+Prosit data sets. Similarly to Fig.8, DIA+Prosit allowed the identification of a higher number of pathways likely involved with plant response to infection.

The proteins identified by DIA that were upregulated in Diseased samples were involved in aminoglycan catabolic process, response to bacterium, cell wall organization or biogenesis, innate immune response, gluconeogenesis, hydrogen peroxide catabolic process, and response to cadmium ion. The most statistically significant pathways (involved in aminoglycan catabolic process, response to bacterium, cell wall organization or biogenesis, and innate immune response) were upregulated in both DIA and DIA+Prosit data. The latter revealed a higher number of proteins thus higher coverage and also showed other pathways such as response to ion and carbon fixation in photosynthetic organisms as more significantly enriched (lower p-values).

The analysis of the downregulated proteins in the Diseased plants showed that except for the arabinan catabolic process, all the other identified pathways were significantly enriched in the DIA+Prosit approach, which revealed galactose metabolism, hexose metabolic process, reductive pentose phosphate cycle and response to cadmium ion.

## 4. Discussion

This was the first DIA study of vascular sap of grapevines using Prosit [33]. We used a pressure chamber to extract the vascular leaf sap from grapevines comparing healthy and diseased plants and submitted samples for proteome analysis. Previous studies of the grapevine xylem proteome have provided important clues regarding the plant responses to infection; however, they have also faced several technical challenges in extracting enough material to adequately describe the complexity of this pathosystem. The focus of this study was to show the application of DIA in combination with Prosit to improve protein prediction and quantification in the vascular sap of leaves infected with *Xylella fastidiosa*. Our results suggest that incorporating a deep learning architecture approach like Prosit to DIA data could help researchers identify more protein candidates in response to pathogenesis and other biological phenomena. Prosit significantly increased the number of proteins, especially in low abundance detected in both from *Vitis* and *Xylella,* contributing to a more detailed picture of this plant-pathogen interaction.

### 4.1. A new proteomic approach for vascular sap studies

The implementation of Prosit to the DIA data increased detected proteins from 145 to 360 for grapevines and from 18 to 90 proteins detected for *Xylella fastidiosa*. Proteomics studies from vascular plant sap have always faced technical challenges due to the low protein concentration present in this plant organ. Previous studies identified differently expressed transcripts and proteins in grapevines by 2D-PAGE for protein isolation and further detection by MS/MS. The maximum resolution for these sample types was around 100 proteins with molecular weights from 20 kDa-75 kDa, with a majority higher than 40 kDa [41]. The most recent proteomic study related to Pierce’s disease detected 91 proteins by LC-MS/MS that ranged from 12 kDa-114 kDa. That study demonstrated that structural data could be incorporated in the pipeline of proteomic data analysis using CHURNER [25]. The number of identified peptides from these 91 proteins also ranged from 2-23 peptides. Combining DIA+Prosit with these complementary functional approaches might provide yet a deeper comprehension of the relevant processes taking place during infection and the molecular functions that could be targeted with priority for increased plant defense.

By using DIA and Prosit, the number of proteins increased as well as the sensitivity of the detection. The number of proteins in low abundance were mostly predicted by Prosit. That’s because the intensity prediction improved the quality of peptide identification by data searching [33]. The molecular weight of proteins from our study ranged from 8 kDa - 217 kDa, significantly broader than in previous studies. The smallest protein predicted by DIA+Prosit was BBE domain-containing protein (VIT_10s0003g05430) with 8 kDa predicted only by DIA+Prosit with six exclusive peptides. This protein has been previously described as necessary in the plant-pathogen interaction of Vitis and *Botrytis cinerea*. BBE-like enzymes inactivate oligo galacturonides (OGs) accumulated as intermediate reaction products of the inhibition of polygalacturonases (PGs) by PG-inhibiting proteins (PGIPs) [42], [43]. By oxidizing OGs, those became less active as defense inducers and less susceptible to hydrolysis of the pathogen’s PGs. The accumulation of OGs can compromise plant growth and resistance through cell death induction. Therefore, the downregulation of BBE-like enzymes in grapevines infected with *Xf* contribute to the plant’s susceptibility. This is the first report of detection of this protein in grape xylem sap, only achieved with Prosit.

The largest protein was a member of the subtilase family (VIT_16s0098g00970), with 217 kDa detected by DIA and DIA+Prosit. These proteins control the establishment of systemic induced resistance and immune priming by the detection of the biotic stimulus [44]. This protein was not detected in Healthy plants in the DIA data, only in the Diseased plants with seven identified peptides. In the DIA+Prosit data, the number of peptides increased and were then detected in Healthy samples as well, but at lower levels compared to the Diseased. Prosit also increased the number of detected peptides to nine. This result exemplifies the increase in sensitivity by implementing Prosit to DIA data.

### 4.2. Plant response to *X. fastidiosa* infection as assessed by the vascular sap

Although the number of studies investigating expressed transcripts and proteins in the xylem sap of plants infected with *Xf* is small, they have provided valuable information regarding plant responses to infection [23], [25], [41]. By accurately evaluating the vascular leaf sap of infected plants with *Xf* using a more sensitive and reproducible proteomic approach, our study confirmed the presence of secreted proteins associated with pathogenesis-related (PR) proteins, chitinases, and β-1-3-glucanases as the key players in mediating the defense response upon pathogen infection [25]. Our study specifically revealed β-1-3-glucanase 3 (VIT_08s0007g06060) as the vital protein contributing to the variance between Healthy and Diseased plants in the DIA+Prosit data (VIP = 3.1) and the second most important in the DIA data (VIP = 2.7). A total of five β-1-3-glucanases proteins were also detected in the DIA+Prosit, and only four were in the DIA data. Except for one protein (VIT_06s0061g00100) that was slightly downregulated in the diseased plants, all the others in both data sets were upregulated. β-1-3-glucanases belong to the PR2 class, and their expression is induced by several pathogens including fungi, oomycetes and most recently shown to be induced by a bacterial infection [25], [41], [45], [46]. Other PR proteins (including PR1), proteases, chitinases, and peroxidases were also confirmed in our study but in a higher number of proteins. Chakraborty et al. (2016) were able to detect 15 peroxidases, and our DIA and DIA+Prosit increased this number to 20. This could be due to the Prosit predictions being generalized to non-tryptic peptides increasing peptide predictions [33].

## 5. Conclusions

This study demonstrated a successful example of using the DIA approach combined with deep learning neural network Prosit for analysis of proteomic data. A total of 360 proteins were identified and quantified from the grapevines subjected to *Xf* inoculation. We also identified different sets of proteins regulated upon infection that were previously shown in other proteomic studies and highlighted new low molecular weight and low abundance proteins previously undetected. This is especially useful in samples with a lower protein abundance and diversity, providing more functional clues of significant players.

**Table 1.** Overview of Proteomics studies of vascular sap of grapevines.

| *Vitis sp.* variety | Biological material | *X. fastidiosa* inoculation | Method | Peptide spectra Analysis | Total proteins | Protein size (kDa) | Matched peptides | Signal peptide detection | Reference |
| --- | --- | --- | --- | --- | --- | --- | --- | --- | --- |
| Chardonnay | Xylem sap | No | 2D-PAGE<br>MALDI-TOF MS/MS | GPM | 10 | 25-150 | 1 | No | Agüero et al., 2008 |
| PD tolerant and susceptible varieties | Xylem sap | No | 2D-PAGE<br>LC-MS/MS | Mascot | ~100 | 20-75 | 1-4 | No | Bascha et al., 2010 |
| PD tolerant and susceptible varieties | Stem | Yes | 2D-PAGE<br>nano-LC-MS/MS | Bioworks | ~200 | 14.4-45 | 2-32 | No | Yang et al., 2010 |
| Chardonnay | Leaf and Apoplastic fluid | No | 2D-PAGE<br>MALDI-TOF MS/MS | Mascot | 227 and 89 | 15-120 | - | No | Delaunois et al., 2013 |
| PD tolerant and susceptible varieties | Xylem tissue | No | 2D-PAGE<br>MALDI-TOF MS/MS | Mascot | ~200 | 20-75 | - | No | Katam et al., 2015 |
| Thompson Seedless | Xylem sap | Yes | LC-MS/MS | Scaffold | 91 | 10-114 | 2-23 | Yes | Chakraborty et al., 2016 |
| Thompson Seedless | Vascular leaf sap | Yes | LC-MS/MS | ScaffoldDIA | 145 | 12-217 | 2-22 | Yes | *This study* |
| Thompson Seedless | Vascular leaf sap | Yes | LC-MS/MS | ScaffoldDIA + Prosit | 360 | 8-217 | 2-31 | Yes | *This study* |

## 6. Acknowledgments and Funding

We thank Samuel Metcalf for sharing his expertise with the pressure chamber and Ken A. Shackel for allowing us to use his equipment. This work was supported by grants obtained from the California Department of Food and Agriculture Pierce’s Disease Board (CDFA-PD Board). LC-MS was supported by a NIH shared instrumentation grant S10OD021801. C.H.D.S was supported by a Coordination for the Improvement of Higher Level Personnel grant (Coordenação de Aperfeiçoamento de Pessoal de Nível Superior Brazil, No. 99999.013202/2013-08).

## 7. Author Contributions

CHDS and AMD conceived and designed the experiments; CHDS coordinated and performed experiments, functional analysis and wrote of the manuscript edited by PAZ and AMD; RABA contributed in the data analysis and discussions; HS helped data analysis using MetaboAnlyst; MS helped with protein digestion and performed LC-MS/MS and chromatogram library creation; AJ helped plant material and inoculations, BSP conducted proteome data analysis, raw data processing and spectral library search; AMD and all others revised the final manuscript.

---

**Figure S1.**
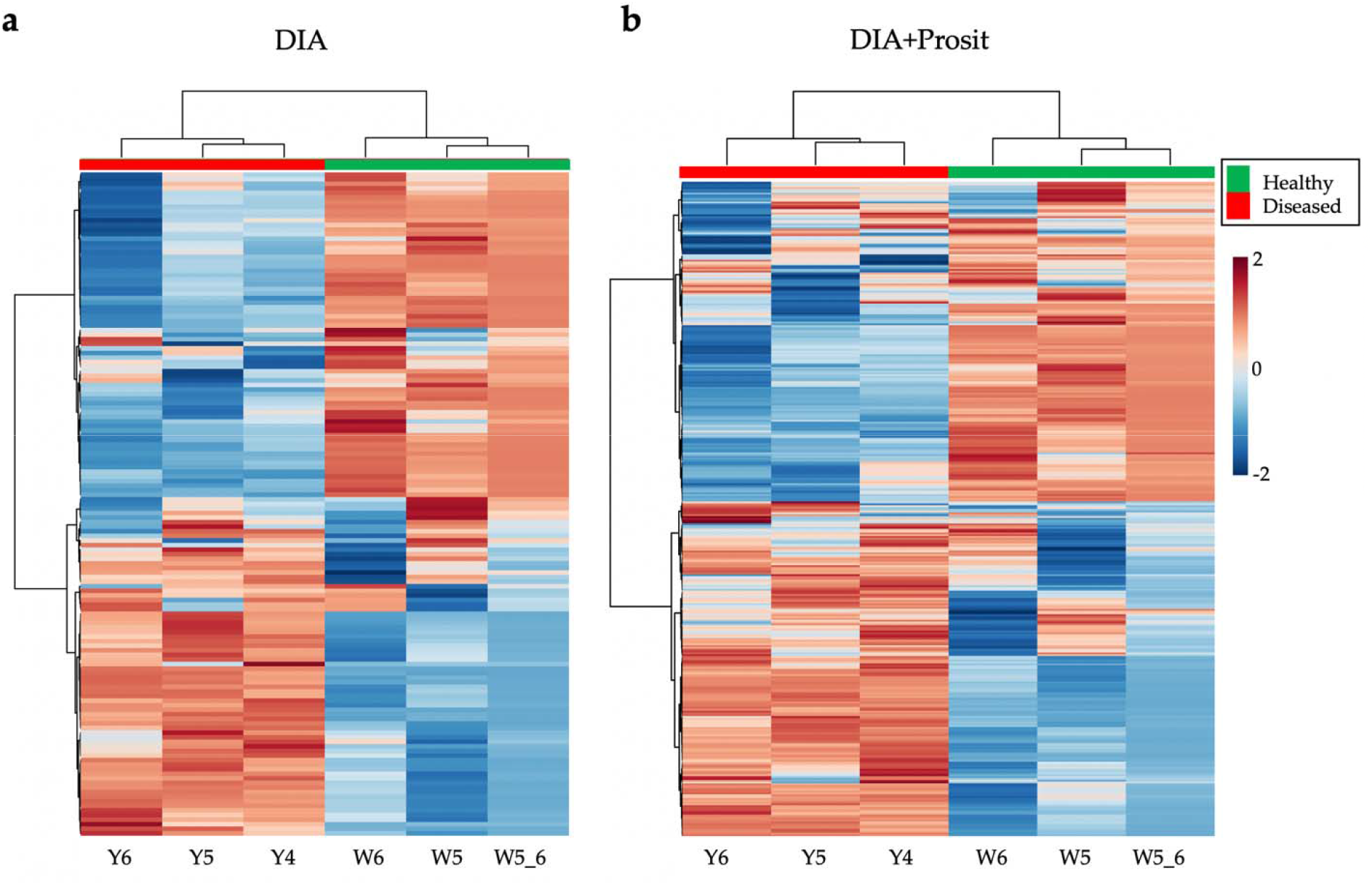
Heat map visualization of the effects of *X. fastidiosa* in grapevines through proteomic analysis of vascular sap of leaves. Hierarchical clustering using Euclidean distance and Ward’s linkage for the clustering algorithm. Samples Y4, Y5, Y6 from Diseased plants and W5 and W6 from Healthy plants. Sample W5_6 is a virtual sample made from the average of W5 and W6. Log_10_ of the exclusive intensity was used from a) DIA and b) DIA+Prosit data sets.

**Figure S2.**
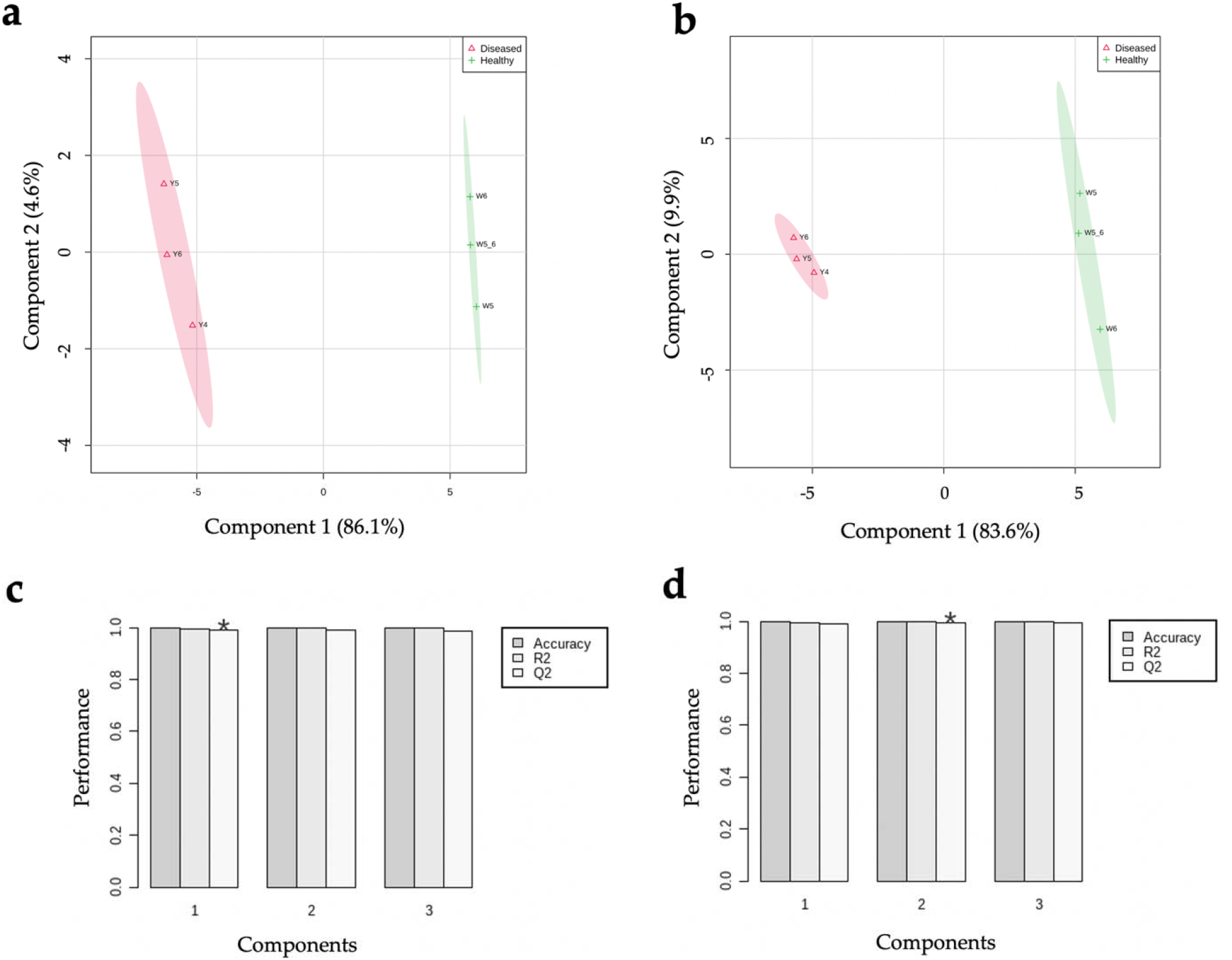
PLS-DA plots and cross-validation of *V. vinifera* samples of Diseased and Healthy plants using a) for DIA data and b) for DIA+Prosit data set. Validation of both models shown by R2 (the sum of squares captured by the model) and Q2 (cross□validation of R2) for the first three components for c) DIA and d) DIA+Prosit. By using Q2, the star indicates the best number of components for the model.

